# *In silico* prediction of high-resolution Hi-C interaction matrices

**DOI:** 10.1101/406322

**Authors:** Shilu Zhang, Deborah Chasman, Sara Knaack, Sushmita Roy

## Abstract

The three-dimensional organization of the genome plays an important role in gene regulation by enabling distal sequence elements to control the expression level of genes hundreds of kilobases away. Hi-C is a powerful genome-wide technique to measure the contact count of pairs of genomic loci needed to study three-dimensional organization. Due to experimental costs high resolution Hi-C datasets are available only for a handful of cell lines. Computational prediction of Hi-C contact counts can offer a scalable and inexpensive approach to examine three-dimensional genome organization across many cellular contexts. Here we present HiC-Reg, a novel approach to predict contact counts from one-dimensional regulatory signals such as epigenetic marks and regulatory protein binding. HiC-Reg exploits the signal from the region spanning two interacting regions and from across multiple cell lines to generalize to new contexts. Using existing feature importance measures and a new matrix factorization based approach, we found CTCF and chromatin marks, especially repressive and elongation marks, as important for predictive performance. Predicted counts from HiC-Reg identify topologically associated domains as well as significant interactions that are enriched for CTCF bi-directional motifs and agree well with interactions identified from complementary long-range interaction assays. Taken together, HiC-Reg provides a powerful framework to generate high-resolution profiles of contact counts that can be used to study individual locus level interactions as well as higher-order organizational units of the genome.

## Introduction

The three-dimensional organization of the genome has emerged as an important component of the gene regulation machinery that enables distal regulatory elements, such as enhancers, to control the expression of genes hundreds of kilobases away. These long-range interactions can have major roles in tissue-specific expression [1–3] and how regulatory sequence variants impact complex phenotypes [4, 5], including diseases such as cancer, diabetes and obesity [6, 7]. Chromosome Conformation Capture (3C) technologies such as, 4C, 5C, ChIA-PET, Hi-C [8] and Capture-Hi-C [9], used to measure 3D proximity of genomic loci have rapidly matured over the past decade. However, there are several challenges for identifying such interactions in diverse cell types and contexts. First, the majority of these experimental technologies have been applied to well-studied cell lines. Second, very few Hi-C datasets are available at resolutions high enough (e.g, 5kbp) to identify enhancer gene interactions due to the tradeoff between measuring genome-wide chromosome organizations while achieving higher resolutions. Third, long-range gene regulation involves a complex interplay of transcription factors, histone marks and architectural proteins [10–13], making it important to examine three-dimensional proximity in concert with these components of the transcription machinery.

Recently, numerous computational approaches for predicting long-range interactions have been developed [14–18], which leverage the fact that regulatory regions that participate in long-range regulatory interactions have characteristic one-dimensional genomic signatures [14, 19]. However, these methods have all used a binary classification framework, which is not optimal because the interaction prediction problem is a one-class problem with no negative examples of interactions. In addition, these do not exploit the genome-wide nature of high-throughput chromosome capture datasets such as Hi-C as they focus on significantly interacting pairs, which can be sensitive to the method used to call significant interactions. Recent comparison of methods for calling significant interactions showed that there are substantial differences in the interactions identified from different methods [20].

To overcome the limitations of a classification approach and to maximally exploit the information in high-throughput assays such Hi-C, we developed a Random Forest regression-based approach, HiC-Reg. HiC-Reg integrates published Hi-C datasets with one-dimensional regulatory genomic datasets such as chromatin marks, architectural and transcription factor proteins, and chromatin accessibility, to predict interaction counts between two genomic loci in a cell line-specific manner. We applied our approach to high-resolution Hi-C data from five cell lines to predict interaction counts at 5kb resolution and systematically evaluated its ability to predict interactions within the same cell line (tested using cross-validation), across different chromosomes as well as different cell lines. Our work shows that a Random Forests-based regression framework can predict genome-wide Hi-C interaction matrices within cell lines and can generalize to new chromosomes and cell lines and modeling the signal between regions as well as integrating data from multiple cell lines is beneficial for improved predictive power. Feature analysis of the predicted interactions suggests that CTCF and chromatin signals are both important for high quality predictions. Computational validation of HiC-Reg shows that HiC-Reg predictions agree well with ChIA-PET datasets, recapitulate well-known examples of long-range interactions, exhibit bidirectional CTCF loops, and can recover topologically associated domains. Overall, HiC-Reg provides a computational approach for predicting the interaction counts discovered by Hi-C technology, which can be used for examining long-range interactions for individual loci as well as for studying the organizational properties of chromosome conformation.

## Results

### HiC-Reg: A Random Forests regression approach for predicting high-resolution contact count of genomic regions

We developed a regression-based approach called HiC-Reg to predict cell line-specific contact counts between pairs of 5kb genomic regions using cell line specific one-dimensional regulatory signals, such as histone marks and transcription factor binding profiles (Fig 1). Our regression framework treats contact counts as outputs of a regression model from input one-dimensional regulatory signals associated with a pair of genomic regions. As cell line specific training count data we used high resolution (5kb) Hi-C datasets from five cell lines from Rao et al [21]. We used features from 14 cell line specific regulatory genomic datasets and genomic distance to represent a pair of regions. These datasets include the architectural protein CTCF, repressive marks (H3k27me3, H3k9me3), marks associated with active gene bodies and elongation (H3k36me3, H4k20me1, H3k79me2), enhancer specific marks (H3k4me1, H3k27ac), activating marks (H3k9ac, H3k4me2, H3k4me3), cohesin component (RAD21), a general transcription factor (TBP) and DNase I (open chromatin) [14]. HiC-Reg uses a Random Forests regression model as its core predictive model. The Random Forests regression model significantly outperforms a linear regression model suggesting that there are non-linear dependencies that are important to be captured to effectively solve the count prediction problem (**Supplementary Fig 1**).

**Fig 1.**
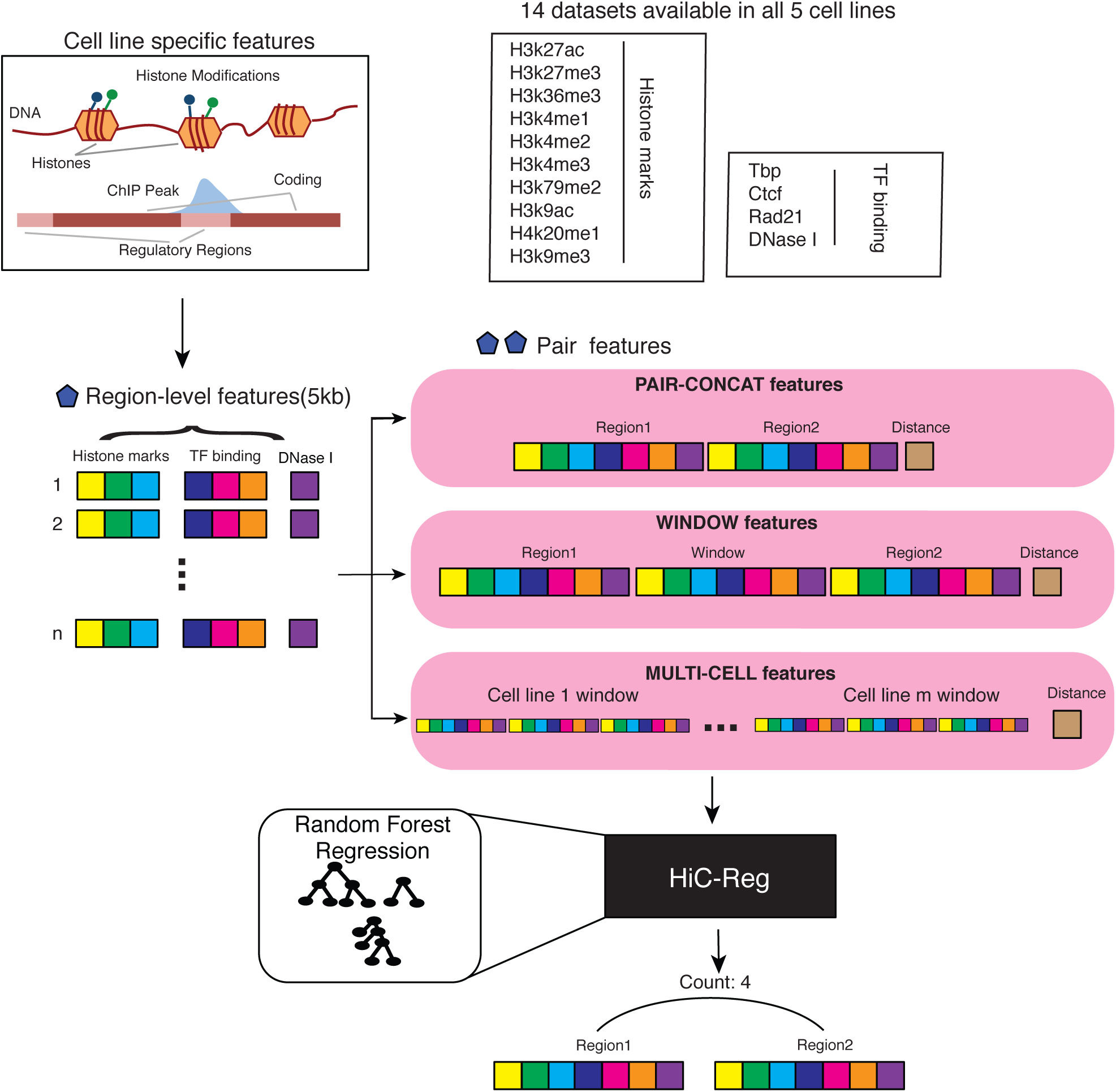
Overview of the HiC-Reg framework. HiC-Reg makes use of 14 datasets: 9 chromatin marks and 5 datasets related to Transcription Factor (TF) binding. A 5kb genomic region is represented by a vector of aggregated signals of either chromatin marks, accessibility, or TF occupancy. A pair of regions in HiC-Reg is represented using one of three types of features: PAIR-CONCAT, WINDOW, and MULTI-CELL. HiC-Reg uses Random Forests regression to predict the contact count between pairs of genomic regions. HiC-Reg takes as input a feature vector for a pair of regions and gives as output a predicted contact count for that pair (e.g., Count:4).

A key consideration for chromosomal count prediction is how to represent the genomic region pairs as examples for training a regression model. To represent a pair in HiC-Reg, we considered three main feature encodings in addition to genomic distance between the two regions of interest: PAIR-CONCAT, WINDOW and MULTI-CELL (Fig 1). PAIR-CONCAT simply concatenates the features for each region into a single vector. The WINDOW feature encoding additionally uses the average signal between two regions and was proposed in TargetFinder, a classification-based method [15]. MULTI-CELL incorporates signals from other cell lines. We assessed the performance of these methods using distance stratified Pearson’s correlation computed on test set pairs in a five fold cross validation setting (**Methods**). The distance stratified correlation measures the correlation between predicted and true counts for genomic pairs at a particular distance threshold. Distance stratification is important due to the high dependence of contact count on genomic distance [22]. As a baseline we compared the performance of these models to a Random Forest regression model trained on distance alone. The performance of HiC-Reg is much better than the performance of distance alone, demonstrating that addition of regulatory signals significantly improves performance (Fig 2). Between PAIR-CONCAT and WINDOW, WINDOW features are signif-icantly better and the performance of PAIR-CONCAT often decreases as function of distance (Fig 2A). WINDOW and MULTI-CELL have similar performance and are both better than using PAIR-CONCAT. To examine the generality of this behavior across all chromosomes, we applied HiC-Reg in all chromosomes in a five fold cross validation setting. To summarize the performance captured in the distance stratified correlation curve, we computed the area under the distance stratified correlation curve (AUC, **Methods**). The AUC metric ranges from 0 to 1, with 1 representing the best performance and succinctly describes the performance across all five cell lines and chromosomes. We find that across cell lines and all chromosomes, both the WINDOW and the MULTI-CELL features are significantly better than simple concatenation of the features of regions suggesting that incorporating the signal between two regions is important for capturing these interaction counts at longer distances (Fig 2B). When examining the performance across the five cell lines, the AUC for the Nhek cell line was lowest suggesting it is the hardest to predict. Due to the superior performance of the WINDOW and MULTI-CELL features, we excluded the PAIR-CONCAT feature from subsequent experiments.

**Fig 2.**
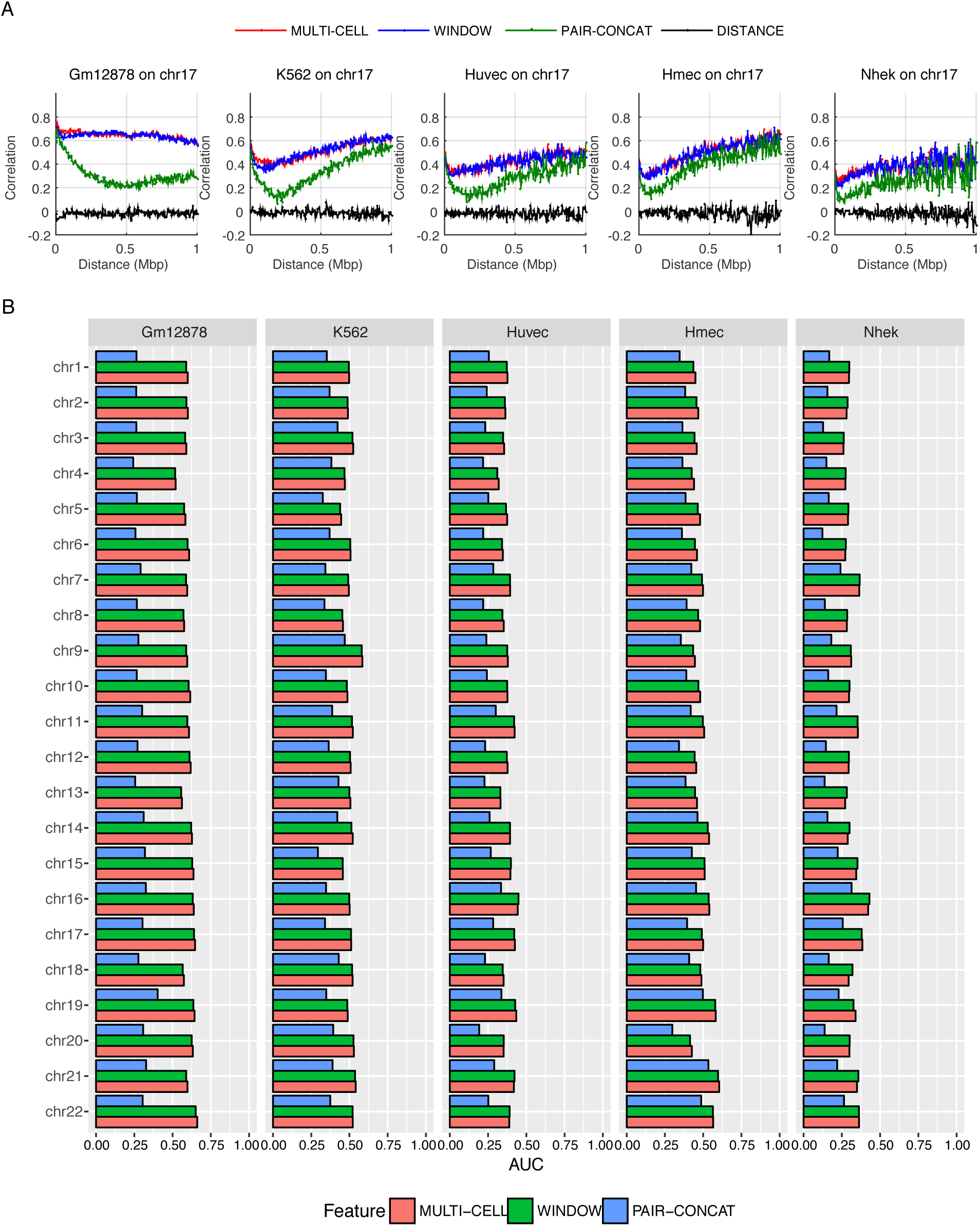
HiC-Reg cross-validation performance. **A.** The distance-stratified Pearson’s correlation plots of test data when training on the same cell line for chromosome 17 in five cell lines: Gm12878, K562, Huvec, Hmec, Nhek. The x-axis corresponds to a particular distance bin and the y-axis corresponds to the Pearson’s correlation of the counts predicted by HiC-Reg for pairs at a particular distance and the true counts. The correlation plots for three types of feature representations of pairs are shown: MULTI-CELL, WINDOW and PAIR-CONCAT. **B.** Area under the curve (AUC) for the distance-stratified Pearson’s correlation in all chromosomes for each of the five cell lines. The AUC is computed for test pairs using 5 fold cross validation.

To examine whether our approach can generalize across chromosomes, we trained HiC-Reg on one chromosome and used it to predict counts in a different chromosome for the same cell line. We considered five chromosomes, chr9, chr11, chr14, chr17 and chr19, each as training and test chromosomes, in all five cell lines. HiC-Reg predictions across chromosomes were slightly worse than when training and testing on the same chromosome in all but the Gm12878 cell line, where the cross chromosome performance was much worse. Interestingly, while both feature types, WINDOW and MULTI-CELL were comparable when training and testing in the same chromosome setting (Fig 2), when comparing models across chromosomes, we found that the MULTI-CELL feature was better than the WINDOW feature (Fig 3, **Supplementary Figs 2-6**), especially for the Gm12878 cell line. Some chromosomes produced worse models than other chromosomes. For example, chromosome 19 was a poor predictor of other chromosomes in all cell lines, and chromosome 14 was a poor predictor of all other chromosomes in K562. These performance differences are indicative of unique chromosomal features that are shared or unique to specific cell lines (chr14 in K562). Taken together, our analysis showed that HiC-Reg is able to successfully predict interaction counts in different cell lines using one-dimensional features and the MULTI-CELL features in particular are important for a cross-chromosome setting.

**Fig 3.**
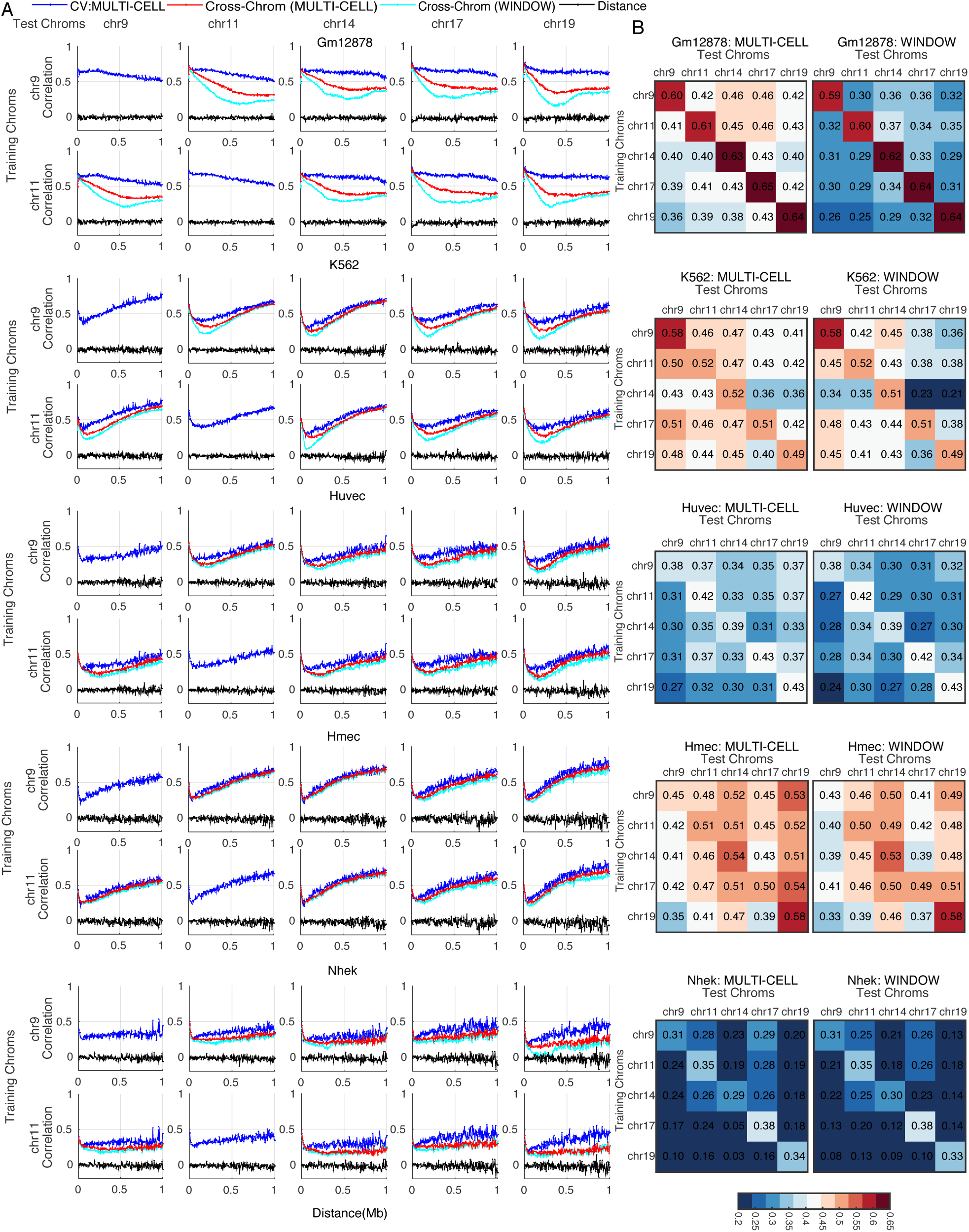
HiC-Reg cross-chromosome performance. **A.** The distance-stratified Pearson’s correlation plot when training on one chromosome and testing on a different chromosome from the same cell line. Shown are the distance-stratified correlations for models trained on chromosome 9 and 11 (row groups) and tested on chromosome 9, 11, 14, 17, 19 (columns) at 5kb in different cell lines. Each pair of rows corresponds to a particular cell line and results are shown for five cell lines: Gm12878, K562, Huvec, Hmec, Nhek.**B.** Heatmap of AUC for all pairs of tested cross-chromosome experiments. Each off-diagonal entry in the heatmap denotes the AUC when trained on the row chromosome and tested on the column chromosome. The more red an entry the better the performance in that chromosome pair combination. The diagonal entries are the AUC values when training and testing on the same chromosome in cross-validation mode.

### Feature analysis identifies important determinants of chromosome contact counts

To gain insight into the relative importance of different one-dimensional regulatory signals such as chromatin marks and transcription factor binding signals for predicting contact counts, we conducted different types of feature analyses on our learned random forests regression models. We focused on the MULTI-CELL features as they had the best performance. We first ranked features using a standard feature importance score, “Out Of Bag” (OOB) error (**Methods**, Fig 4A, **Supplementary Fig 7**). For a given cell line, the features from the cell line ranked highly (e.g., in Gm12878 all top 10 features were from the Gm12878 cell line, Fig 4A); however, there were some exceptions where a feature from another cell line was also important (e.g., several Gm12878 features were important for the HUVEC and K562 cell lines, Fig 4A). The feature rankings across different chromosomes were similar (Spearman correlation 0.72-0.87, **Supplementary Fig 7D**). Based on this ranking, the most important features included Distance, elongation chromatin marks like H3K36me3 and H4K20me1, a repressive mark H3K9me3, enhancer associated mark H3K4me1, and DNase I, typically on one or both end point regions (R1 and R2) and rarely on the Window regions (W). We next used a complementary strategy of counting the number of times a feature was used for predicting the count of a test pair (Fig 4B, **Supplementary Fig 8**). This analysis also found Distance to be important but also other factors like CTCF, TBP, DNase I, as well as repressive marks (H3k9me3 and H3k27me3), an enhancer associated mark (H3K4me1), and an elongation mark (H4K20me1). Here too we observed that the features measured in the training cell line were typically most highly ranked with some variations (e.g., Gm12878 features were useful for predicting counts in most cell lines). Comparing across chromosomes, the feature rankings were very similar (Spearman correlation 0.85-0.94 **Supplementary Fig 8D**). The two feature analysis methods agreed on the importance of elongation mark, H4K20me1, enhancer mark H3K4me1 and several repressive marks, but there were some differences. The OOB method identified histone mark features in the region (R1 and R2) while the feature usage counting importance implicated CTCF and TBP Window signals to be the most important.

**Fig 4.**
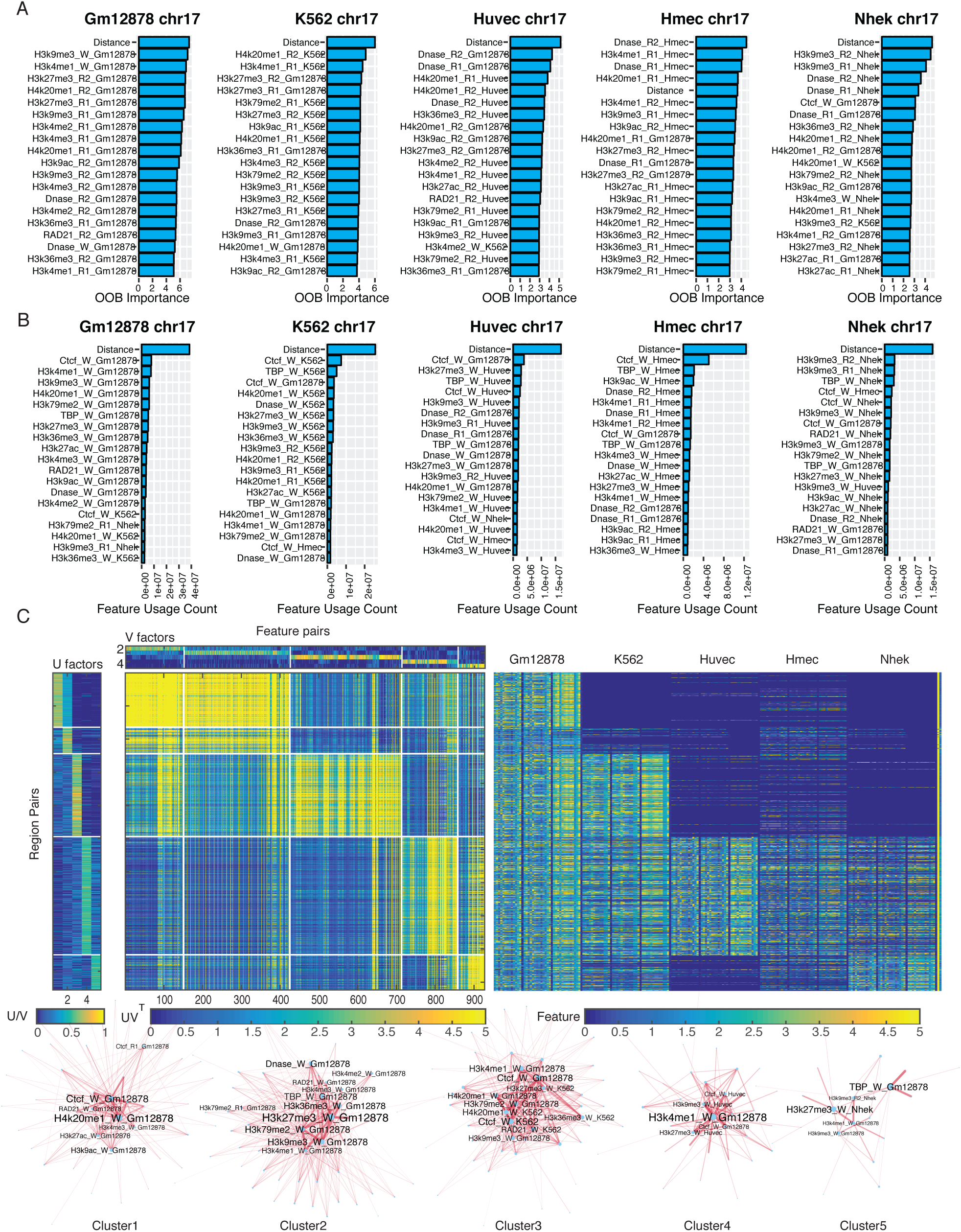
Analysis of features important for predicting Hi-C contact counts. **A.** Shown are the top 20 MULTICELL features ranked based on Out Of Bag (OOB) feature importance on chromosome 17 in all five cell lines. Each horizontal bar corresponds to one feature. The feature name includes the name of the histone mark, DNase I or TF, whether it is on one of the interaction regions (R1, R2) or in the intervening window (W), and the specific cell line from which this feature is extracted. **B.** Shown are top 20 features ranked based on counting the number of times a feature is used for test set predictions. Feature rankings are for chromosome 17 for all five cell lines. **C.** Non-negative matrix (NMF) factorization of region-pair by featurepair matrix for Gm12878 chromosome 17. The *U* and *V* factors are the NMF factors to provide membership of region pairs or feature pairs in a cluster (white lines demarcate the region pair and feature pair clusters). The factorized feature count matrix is shown below the *V* factors and to the right of the *U* factors. The heatmap on the right are the features associated with each of the pairs, with rows corresponding to a pair of regions and columns corresponding to the feature values grouped by the cell line from which they are obtained. Bottom are Cytoscape network representation of important pairs of features. The node size is proportional to the number of features the specific feature co-occurs on a path in the decision tree. The thickness of the line is proportional to the number of times the pair of features is used on the path from root to the leaf for a test example pair. Font size of the node label is proportional to its size.

**Fig 7.**
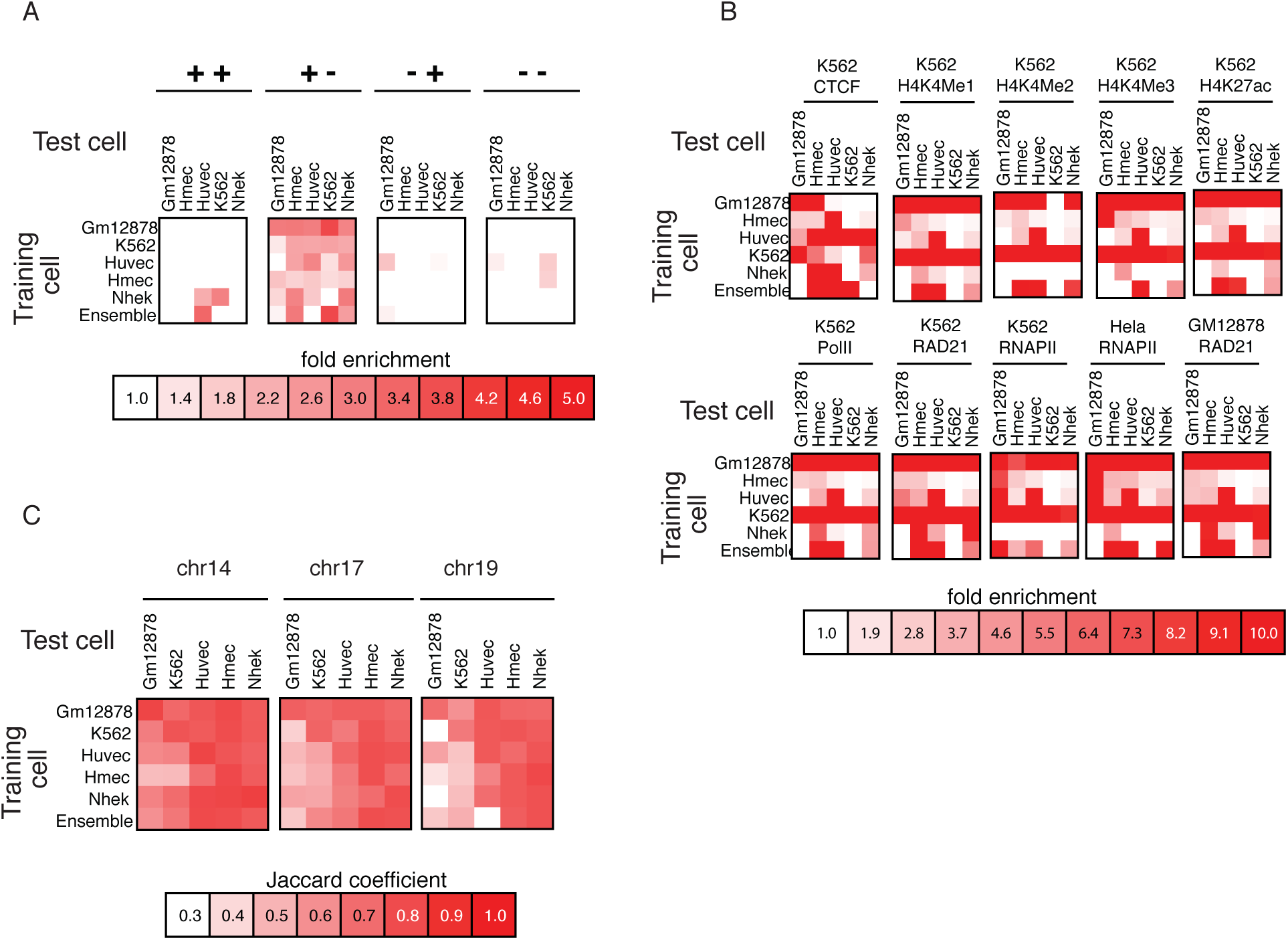
Ability of HiC-Reg to capture significant interactions and TADs in new cell lines. **A.** Enrichment of CTCF motifs in four configurations for cross-cell predictions. Shown are the fold enrichments when testing using a model trained on different cell line or an ensemble of models using MULTI-CELL features. The diagonal entries correspond to the fold enrichment obtained in the cross-validation setting. **B.** Fold enrichment of ChIA-PET interactions in predicted interactions when trained on a different cell line as well as when using the MULTI-CELL ensemble. **C.** Jaccard coefficient based similarity of TADs called on true and predicted interaction counts when training on a different cell line. Shown are Jaccard similarity values for three chromosomes, 14, 17 and 19.

**Fig 8.**
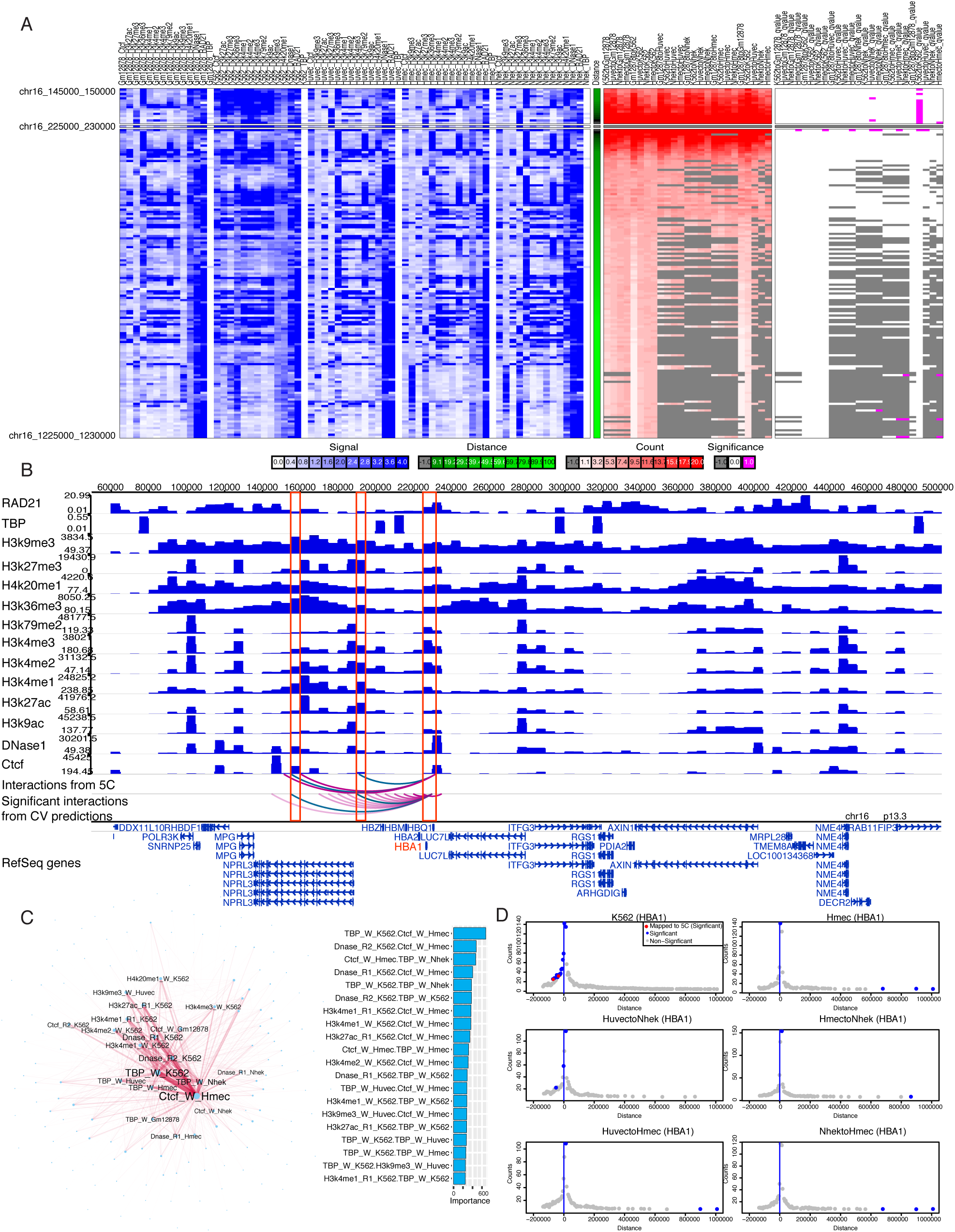
Examining HiC-Reg predictions at the HBA1 locus. **A.** Shown are the features, predicted counts and P-value for the 1MB radius around the 5kb bin spanning the HBA1 gene TSS. The predicted counts (white read heat map) and p-value significance (white magenta heatmap) for the 5 CV models and the 20 cross-cell line models are shown. The columns correspond to the specific model. Gray corresponds to missing count data in the original Hi-C dataset. **B.** Visualization of feature signals using WashU Epigenome Browser for significant interactions obtained from a model trained in K562 and tested in K562. Shown are also overlapping interactions with 5C pairs. Red vertical lines demarcate the 5kb bin overlapping the HBA1 gene TSS and the distal region that interacts with it, as supported by 5C. The green arcs depict the interactions predicted by HiC-Reg and supported by 5C. **C.** Pairwise feature analysis based on pairs of features used for predicting significant interactions associated with HBA1 gene in the K562 cell line (Left: Network representation of important pair-wise features. Right: Ranking of top 20 important pair-wise features). **D.** Manhattan style plots of predicted counts around the 1MB radius of the HBA1 gene. Only predictions from models that had significant interactions are shown (6 out of 25). The left part of each Manhattan plot is shortened because this region is towards the beginning of the chromosome. Blue dots are significant interactions and red dots are significant interactions that overlap a 5C interaction.

While the above approach identified the top features for all pairs of regions, it does not tell us whether there are different feature sets useful for different sets of pairs. In particular, there could be some interacting pairs that are driven largely by chromatin marks, while another set of pairs driven by transcription factors. Furthermore, it does not inform us about dependencies among features that might be important for making these predictions. To address this, we developed a novel feature analysis method, based on non-negative matrix factorization (NMF, **Methods**). Briefly, we obtained region pairs that were in the lowest 5% of test error and counted the number of times a feature or a pair of features was used on the tree path traversed for these region pairs. We selected only feature pairs that were in the 5% lowest errors for computational efficiency and because the individual feature rankings did not change compared to when considering all pairs (**Supplementary Fig 9, 10**). Furthermore, we focused on chromosome 17 because the feature importances were similar across chromosomes (**Supplementary Fig 7, 8**). We used these counts to create a region-pair by feature-pair matrix, with each entry of the matrix denoting the number of times the feature pair was used in the trees. We next applied NMF on this matrix to obtain clusters of region pairs associated with clusters of feature pairs (Fig 4C). Such “bi-clusters” are indicative of different classes of region pairs and the most important features associated with them (Fig 4C). For example, for Gm12878, one cluster (Cluster 1) was associated with H4K20me1 and CTCF, while another cluster (Cluster 2) was associated with repressive marks (H3K27me3) together with other histone marks but not CTCF. A third cluster (Cluster 3) was associated with features from the K562 cell line (CTCF and H4k20me1) in addition to those from Gm12878. Cluster 4 was associated with CTCF in Gm12878 and Huvec and H3K4me1 in Gm12878 and finally Cluster 5 was largely associated with TBP. We found similar behavior in other cell lines as well (**Supplementary Fig 11-14**). For example, in K562, we found two types of clusters: one cluster (Cluster 2), was associated with different chromatin marks, while the other clusters had CTCF, in combination with different chromatin marks. While CTCF was important as a feature hub in all cell lines, some lines exhibited importance of other types of features like H3k9me3 (Nhek), DNase I (Huvec and Hmec) and RAD21 (Nhek). We applied the same procedure to the matrix of region-pairs by individual features (**Supplementary Fig 15, 16**), however, the groups were essentially driven by the cell line in which the feature was measured. While this showed that NMF captures the most important grouping structure (cell line), it was not unexpected and only served as a sanity check.

Taken together, our feature analysis showed that there were largely CTCF-driven and chromatin mark driven clusters of interactions. CTCF was a key feature for predicting interactions and was associated with other chromatin marks. Furthermore, we found that elongation marks like H3K36me3 or H4K20me1 and repressive marks like H3K27me3 were often important as feature hubs.

### HiC-Reg predictions exhibit hallmarks of true looping interactions and identify TADs

We next assessed the quality of our predictions based on several additional metrics that examined the occurrence of specific properties of true interactions. One such property is the occurrence of bi-directional CTCF motifs [21]. Briefly, a pair can have one of four configurations of the CTCF motif, (++), (+-), (-+) and (–), where “+” corresponds to the motif on the forward strand and “-” corresponds to the motif being on the reverse strand. The pairs with the (+-) configuration are most likely the true loops. This property of looping interactions was also used by Forcato et al [20] to compare different Hi-C peak calling programs. To assess the occurrence of CTCF bidirectional motifs in HiC-Reg predictions, we applied Fit-Hi-C [22], to identify significant interactions in both true and predicted counts (**Methods**). Following [20], we only focus on pairs with at least one CTCF signed motif mapped to each pair of regions, but discarding a pair if one of the regions has both orientations of the motif in the same region. We quantified the tendency of CTCF bidirectional motifs to occur in the significant pairs versus all pairs using fold enrichment, which compares the fraction of interactions predicted by HiC-Reg that has a particular configuration compared to the proportion of interactions expected by random chance. Across all five cell lines, significant pairs called on both the true and predicted counts are enriched for the bidirectional motif (+-) configuration (Fig 5A). The level of enrichment is comparable for the interactions identified from the true and predicted counts and is sometimes better in the predicted counts.

**Fig 5.**
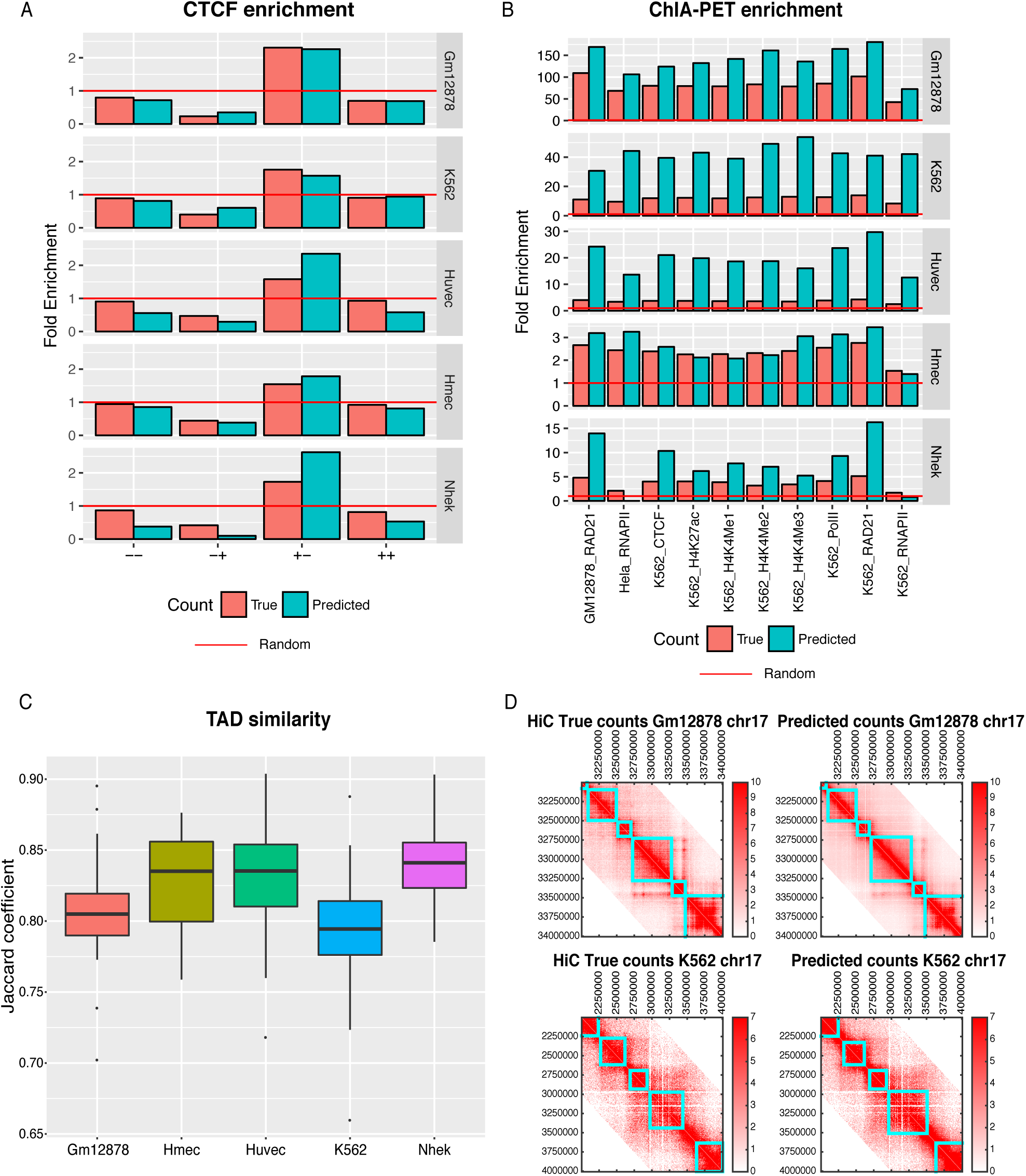
Assessing significant interactions and TADs from HiC-Reg predictions. **A.** Fold enrichment of four configurations of CTCF motifs in significant interactions identified using true and predicted counts. A fold enrichment >1 (red horizontal line) is considered as enriched. Fold enrichment in all five cell lines is shown. **B.**Fold enrichment of interactions identified using ChIA-PET experiments in significant interactions from HiC-Reg’s predictions and true counts. A fold enrichment >1 (red horizontal line) is considered as enriched.**C.**Distribution of TAD similarity identified from true and predicted counts for all chromosomes. Each point in the box plot corresponds to the average Jaccard coefficient for a chromosome. **D.** TADs identified on true (left) and predicted (right) HiC count matrices for selected regions. Top: Gm12878 cell line, chr17:3234Mbp. Bottom: K562 cell line, chr17: 2-4Mbp)

As a second evaluation metric, we compared the significant interactions identified by Fit-Hi-C on HiC-Reg’s predictions and true counts with interactions identified using a complementary experiment, ChIA-PET (**Methods**). We obtained 10 published datasets for different factors (RNA PolII, CTCF and RAD21) and histone marks in multiple cell lines [23, 24]. We estimated fold enrichment of the ChIAPET interactions in the significant interactions compared to background. We found that interactions from both true counts and HiC-Reg predictions were enriched for ChIA-PET interactions, and these enrichments were often better for HiC-Reg predictions than those from the true counts (Fig 5B). The CTCF directionality and ChIA-PET enrichment suggest that significant interactions from HiC-Reg predictions exhibit hallmarks of true looping interactions.

As a third validation metric, we asked to what extent the HiC-Reg counts can be used to study structural units of organization of the chromosome, such as topologically associated domains (TADs, [25]). One of the advantages of a regression versus a classification framework is that the output counts can be examined with downstream TAD finding algorithms [20, 26]. We applied the Directionality Index (DI) TAD finding method to HiC-Reg predicted counts and true counts and compared the similarity of the identified TADs using a metric derived from the Jaccard coefficient (See **Methods**). The Jaccard coefficient assesses the overlap between two sets of objects (e.g., regions in one TAD versus regions in another TAD) and is a number between 0 and 1, with 0 representing no overlap and 1 representing perfect overlap. We aggregated the Jaccard coefficient across all TADs identified on a chromosome into a single average Jaccard coefficient (**Methods**). Across all cell lines and chromosomes, the average Jaccard coefficient ranged between 0.79-0.83 indicating good agreement between TADs from true and predicted counts (Fig 5C). This high agreement is visually shown for a selected region (2Mb block of 5kb regions on chr17: 32000000-34000000) where the identified TADs (cyan boxes) agree between the true and predicted count matrices (Fig 5D).

Overall, these validation results show that HiC-Reg predictions can be used to study three-dimensional genome organization at the level of individual loops as well as the level of higher-order structural units such as topologically associated domains.

### HiC-Reg can be used predict contact counts in new cell lines

Our analysis so far demonstrates the feasibility of using regression to predict chromosome contact count in cell lines with available Hi-C data for at least some chromosomes. We next asked if we could apply this approach to predict interactions in test cell lines different from the training cell line. This would enable us to study the utility of HiC-Reg in cell lines where Hi-C data are not available. We applied HiC-Reg trained on one cell line to predict counts for pairs from a different test cell line. We evaluated the quality of the predictions in the test cell line using the distance-stratified Pearson’s correlation (CrossCell, Fig 6), and additional validation metrics (CTCF directionality, ChIA-PET and TAD recovery, Fig 7). These validation metrics were computed on the test cell line and compared against different models: (i) model trained on distance alone, (ii) model trained with cross-validation (CV) on the test cell line (CV, Fig 6), and (iii) a new baseline model which simply “transferred” the count from the training cell line to the test cell line (TransferCount, Fig 6).

**Fig 6.**
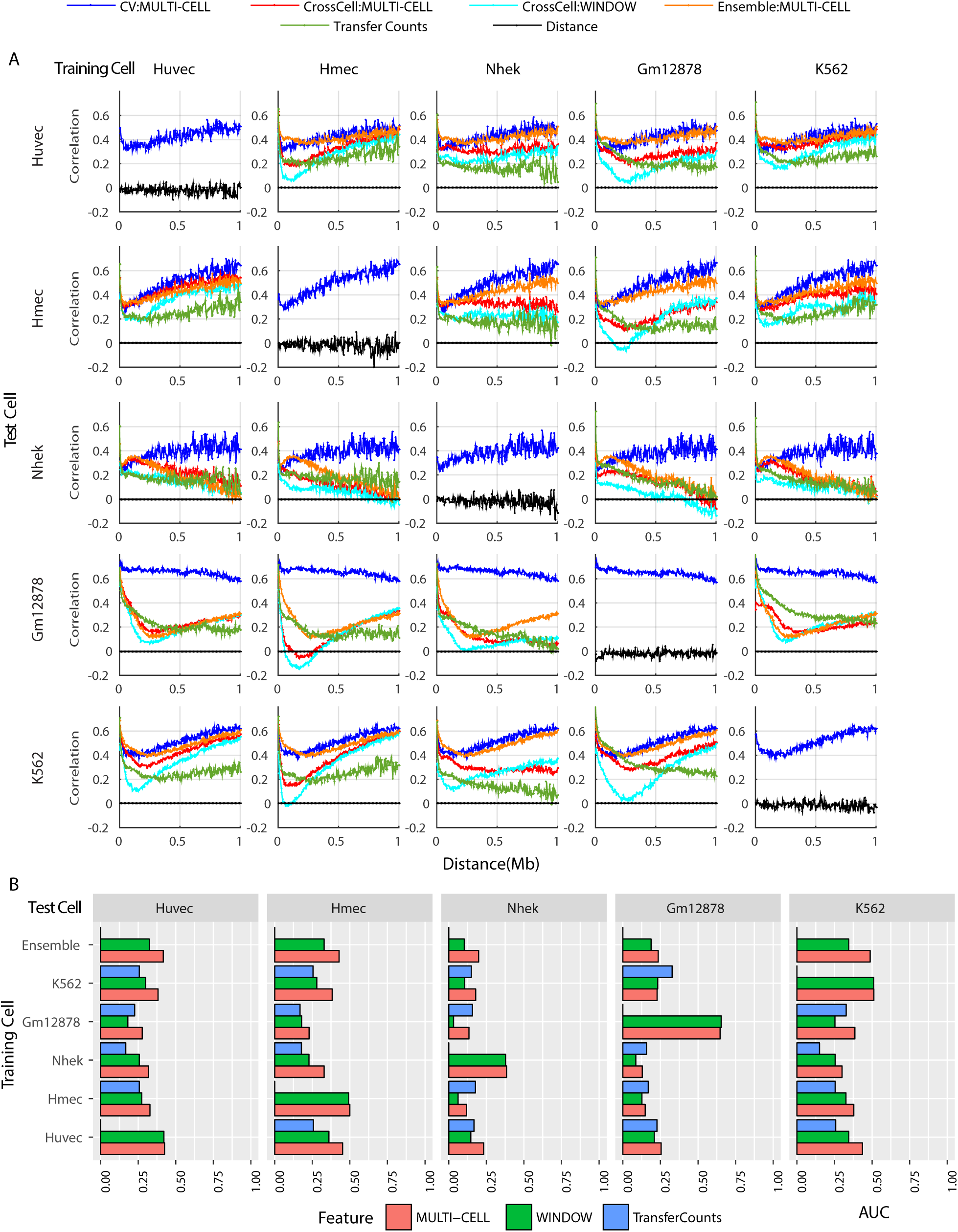
Assessing HiC-Reg predicted counts in new test cell lines. **A.** Distance stratified Pearson’s correlation plot using models trained on one cell line (columns) and tested on a different cell line (rows) for chromosome 17 and all five cell lines: Huvec, Hmec, Nhek, Gm12878, K562. Each plot (except the ones on the diagonal) show the distance stratified correlation plots when using the same cell line (blue), different cell line using MULTI-CELL (red) and WINDOW (cyan) features, using the ensemble of all predictions from a different cell line (orange), when simply transferring counts (green). The plots on the diagonal show the CV performance. **B.** Area under the curve (AUC) for distance stratified correlation in chromosome 17. Shown are the AUCs for the different training cell line models for both feature types (red and green), ensembles from both feature types, as well as transfer count when predicting in a new cell line.

A model trained on a cell line different from the test cell line is significantly better than a model trained on distance alone, but is often worse than a model trained on the same cell line (Fig 6). For example, for chr17, the same cell line CV model has the best performance (blue line Fig 6A) compared to all other versions of cross cell line predictions. Here too we observe that the MULTI-CELL features (Fig 6A, red line) are better or at least as good as the WINDOW features (Fig 6A, cyan line). Compared to transferring counts (TransferCount, green line Fig 6A), both MULTI-CELL and WINDOW have significant benefits at long distance relationships (green line is usually below the red and cyan lines after ~250kb). The summarized AUC offer a concise summary of this behavior (Fig 6B), with models using MULTI-CELL being at least as good as TransferCount models in the majority of training-test cell line combinations. Overall, the Gm12878 cell line was the hardest to predict using a model from other cell lines.

When comparing the predictions in other chromosomes (**Supplementary Fig 17, 18**), we observe a similar behavior. The MULTI-CELL was at least as good or better than transferring counts or WINDOW features and the Gm12878 cell line remained the hardest cell line to predict. Interestingly, the extent to which a cell line can be predicted from a model in a different cell line depends greatly on the test cell line. In particular, on Gm12878 which has the highest sequencing depth, none of the models were able to come up to par with the model trained and tested on Gm12878 (Fig 6A, fourth row, Fig 6B, fourth column). In contrast, for HUVEC, the K562 model is able to predict interactions nearly as well as the CV model especially when using the MULTI-CELL features. Similarly, for HMEC, HUVEC-trained model was able to recapitulate the performance of the HMEC CV-trained model to a great extent (Fig 6B). Previously, we have shown that an ensemble model of combining predictions from multiple models provided a robust performance in a new cell line [14]. Therefore, we combined the predictions from the models trained on each of the cell lines by taking the average of predictions (Fig 6, ENSEMBLE). We find that the ensemble predictions are at least as good as the predictions from the individual models (Fig 6A orange line) and the ensemble for the MULTI-CELL features is better than that for the WINDOW features (Fig 6B). In K562, Huvec and Nhek cell lines, we found that ensemble approaches performed comparably with the cross-validation performance and better than the best performing cross-cell line prediction.

As additional validation of our predictions we tested the significant interactions from these predicted counts for enrichment of ChIA-PET interactions, CTCF bidirectional loops and TAD recovery (Fig 7). We again applied Fit-Hi-C to the true and predicted counts and found that the predicted interactions in each of the chromosomes tested were significantly enriched for bi-directional motifs and ChIA-PET interactions at levels comparable to that from true counts. Furthermore, there was good agreement with the TADs identified in each of these chromosomes based on the Jaccard coefficient score.

Overall, these results suggest that HiC-Reg can be used to predict interactions in new cell lines and it performs better than baseline approaches based on distance alone or simply transferring counts. There is a dependence on the training cell line and an ensemble approach offers a robust way to combine predictions from multiple training models.

### HiC-Reg can recover interactions in manually curated high confidence interaction pairs

So far our results show that HiC-Reg can predict genome-wide Hi-C counts and these counts can be used for identifying significantly interacting pairs as well as higher-order organizational units. To gain deeper insight into the features that drive interactions between two specific loci, we next focused on well-characterized long-range interactions. In particular, we examined two loci, one associated with the hemoglobin gene, HBA1 (Fig 8), and the other one associated with the pregnancy associated plasma A (PAPPA) gene implicated in age-related breast cancer susceptibility.

Distal regulation of the HBA1 gene by a regulatory element 33-48kb away [27] has been experimentally characterized using low throughput [27] and high throughput methods such as 5C [28]. We applied Fit-Hi-C on our predicted counts to obtain significant interactions associated with the 5kb bins containing the HBA1 promoter. We found 14 significantly interacting pairs associated with the HBA1 gene, the majority of which came from the K562 cell line (CV), which is consistent with this gene being specific to erythroid cells [27]. Among the 14 significantly interacting pairs, 2 overlapped with 5C datasets (Fig 8B, D, green lines) at 35kb and 70kb. Visualization of the regulatory signals spanning the HBA1 gene and its interacting regions in the WashU genome browser [29], showed chromatin marks including H3K9me3, H3K20me1, H3K36me3, H3K4me1, H3K4me2, H3K27ac as important features and CTCF and DNase I as additional contributors to these interactions (Fig 8B). Examination of individual and pairwise features based on their usage count in the significant interactions showed that the top features for making these predictions came from the Window region of K562 or HMEC (**Supplementary Fig 19A, B**, Fig 8C), and included features such as TBP, CTCF, DNase I as well as chromatin marks such as H3K4me1 and H3K27ac.

We next investigated the PAPPA gene locus, which is implicated in the development of mammary glands and is a gene of interest in breast cancer studies [30, 31]. The rat ortholog of the PAPPA gene was shown to be regulated by a 8.5kb genomic region called the temporal control element (TCE) in rat mammary epithelial cells [31]. The TCE resides within the MCS5C genomic locus associated with breast cancer susceptibility and is conserved in human and mouse [31]. We examined significant interactions associated with the human PAPPA TSS and found the largest number of significant interactions in the HMEC cell line or in cross cell line predictions using the HMEC model (**Supplementary Fig 20D**). The HMEC cell line, is a primary mammary epithelial cell line, which indicates that these interactions are relevant to the breast tissue. We focus on the significant interactions identified in HMEC when trained in CV mode as these are likely the highest quality. We found a total of 7 significant interactions of which 2 overlap the TCE element. Visualization of the signals (**Supplementary Fig 20B**) and feature analysis on the significant interactions indicated that CTCF, TBP and H3K4me1 (**Supplementary Fig 20C**) measured in the HMEC cell line are important for this interaction. In summary, HiC-Reg predictions provide computational support for the long-range regulation of the PAPPA gene in a relevant human cell line, which was originally studied in the rat mammary cells.

Taken together, our fine-grained analysis of two loci known to be involved in long-range regulatory interactions provided further support of our predictions, highlighted potentially important features that facilitate these interactions and serve as case studies of how HiC-Reg could be used to characterize a particular locus of interest. In both cases we found additional loci that are predicted to interact with these genes, which can be followed with future experiments.

## Discussion

The three-dimensional organization of the genome can affect the transcriptional status of a single gene locus as well as larger chromosomal domains, which can both have significant downstream consequences on complex phenotypes. Although high-throughput chromosome capture conformation assays are rapidly evolving, measuring cell line-specific interactions on a genome-wide scale and at high resolution is a significant challenge. In this work, we described a novel computational approach, HiC-Reg that can predict the contact count of two genomic regions from their one-dimensional regulatory signals, which are available for a large number of cell lines and experimentally more tractable to generate than Hi-C datasets. Because HiC-Reg directly predicts counts, instead of classifying interactions from noninteractions as has been commonly done [14, 15, 19], the output from HiC-Reg can be used to identify significant interactions using peak-calling algorithms (e.g., Fit-Hi-C [22]) as well as examine more large scale organizational properties using domain finding algorithms (e.g, Directionality Index method, [32]).

We studied several technical issues within the HiC-Reg regression framework relevant to model design, feature representation and important datasets for predicting counts. A key challenge we tried to address using HiC-Reg was to generate high resolution interaction counts in a new chromosome or cell line of interest. The former is relevant to predict interactions among regions that might have not have been experimentally assayed. The latter is useful because some cell types and developmental stages may not be amenable to large scale high-throughput 3C experiments and computational predictions could prioritize specific regions for targeted experimental studies. Our cross-chromosome experiments showed that the performance is decreased when training on one chromosome and testing on another. The features identified as important across different chromosomes are very similar (**Supplementary Fig 7, 8**), suggesting that the overall properties governing chromosomal contact are similar across chromosomes, however, there may be fine-grained differences that are not being captured by the Random Forests regression model. Incorporation of additional measurements from transcription factor binding could be beneficial for capturing these differences. Our cross-cell line prediction shows that a regressor trained on one cell line can predict interactions in other cell lines. However, the performance can vary from one training cell line to another, making the choice of the training cell line non-trivial. Ensemble approaches that aggregate predictions from multiple predictive models are a natural way to address this problem. While the simple ensemble predictor was better or comparable to the best performing cell line, more systematic approaches to combine shared information across different cell lines such as considering different weighting strategies will be an important direction of future work.

A second issue we considered was determining the most important genomic datasets for predicting contact counts. Our predictive framework enabled us to study known players of long-range gene regulation such as CTCF and cohesin [33], together with additional components of the transcription machinery such as chromatin marks and general transcription factors, several of which have not been thoroughly characterized in the context of long-range interactions. We examined the importance of these features globally for all pairs, for sets of pairs, and for individual pairs. Our analysis showed that in addition to CTCF, elongation and repressive marks can also be important for predicting counts. The importance of elongation marks such as H3K36me3 in predicting contact count is consistent with existing work [9, 34], which showed that H3K36me3 and elongation-related signals are enriched in regions participating in long-range interactions [9] and higher-order genome organization [34]. The identification of repressive marks such as H3K27me3 could be because of their association in large-scale transcriptional units such as compartments [11], TADs [12] and intra-TAD loops [35], and the specific pairs associated with these repressive marks could be specifically transcriptionally silenced or be in a poised state [11].

We evaluated the predictions from HiC-Reg using different validation metrics, globally using complementary assays, as well as, at specific loci that have been studied in the literature through high quality, albeit low throughput experiments. This was important because predictive performance based on the ability to predict contact counts may not necessarily reveal biological insights. In particular, we compared the statistically significant interactions identified from HiC-Reg with those measured experimentally using ChIA-PET and whether they were enriched for CTCF bidirectional loops. Using both metrics, we showed that HiC-Reg predictions can be used to identify significant interactions that have as good experimental support as those from true counts. We also assessed the ability of HiC-Reg predictions to recover higher order units of organization such as TADs and found good agreement between TADs identified using our predicted counts compared to true counts. We demonstrated the utility of HiC-Reg in studying the long-range regulatory landscape of two loci including the well-studied HBA1 locus as well as a relatively less studied locus, PAPPA. Interestingly, our analysis of the PAPPA locus provided support of long-range regulation of PAPPA, originally identified in rat, in a relevant human cell line.

HiC-Reg exploits the widely available chromatin mark signals that are experimentally easier to measure compared to the Hi-C experiment. HiC-Reg however relies on the availability of these marks in new contexts, which may not always be available. Several groups have started to explore imputation strategies of chromatin marks [36, 37]. Another important direction of future work would be to examine how HiC-Reg performs with imputed marks as this would greatly increase the impact of a predictive modeling framework such as HiC-Reg.

In summary, we have developed a regression-based framework to predict interactions between pairs of regions across multiple cell lines by integrating published Hi-C datasets with one-dimensional regulatory genomic datasets. The predictions from HiC-Reg can be used to identify significant interactions as well as examine higher-order organizational units. As additional chromatin mark signals and Hi-C data become available, our method can take advantage of these datasets to learn better predictive models. This can be helpful to systematically link genes to enhancers as well as to interpret regulatory variants across diverse cell types and diseases.

## Material and Methods

### Learning a cell line-specific regression model for predicting Hi-C interactions matrices

HiC-Reg is based on a regression model to predict contact counts measured in a Hi-C experiment using features derived from various regulatory genomic data sets (e.g. ChIP-seq data sets for histone modifications, transcription factor occupancies, Fig 1). HiC-Reg uses Random Forests as its main predictive algorithm. Random Forests are powerful tree ensemble learning approaches that have been shown to have very good generalization performance [38], and have been applied to a variety of predictive problems in gene regulation [39–41]. We trained Random Forests on the above described different feature encodings and Hi-C SQRTVC normalized contact counts downloaded from Rao et al paper [21]. We experimented with different number of trees (**Supplementary Fig 1D**) and found that beyond 20 trees there was no improvement in performance. Hence we performed all subsequent experiments with 20 trees. We compared the Random Forests regression model to a linear regression model using the WINDOW feature encodings of a pair, which includes the signals associated with each region in the pair as well as the average signal between the two regions, and the Distance between two regions. We found that the non-linear regression model based on Random Forests performs significantly better than a linear regression approach (**Supplementary Fig 1**). We next describe the training and test generation and different feature representations of a pair of regions.

### Generation of training and test sets

To generate training and test datasets for our regression models, we first binned each chromosome into 5kb non-overlapping regions. We randomized the regions and split them into five sets. Each time, we select one of the five sets as the test set of regions and the remaining four as training, repeating this for all five sets of regions. Within each training or test set of regions we generate all pairs of interactions that are within a 1MB radius. Because the Hi-C matrix is symmetric, we need to only predict the upper triangle of the matrix and hence our pairs are not redundant. In each pair, the region with the smaller coordinate is referred to as the “R1” region and the region with the larger coordinate as the “R2” region. We conducted three different types experiments to evaluate the performance of HiC-Reg: **(i)** same cell line, same chromosome cross-validation (CV), **(ii)** same cell line cross chromosome comparison, and,**(iii)** different cell line same chromosome comparison.

For the same cell line same chromosome setting, HiC-Reg was trained and tested using five-fold cross-validation using training and test pairs generated as described above. In each fold, we trained random forests regression models on 4 folds, and predict contact counts for the left-out fold. We concatenated predictions from five folds and assessed performance using distance-stratified Pearson’s correlation of true and predicted counts. Because each training/test example is a pair of regions, we need to consider two types of examples: those that share a region with the training data (easy examples) and those that do not share a region with the training data (hard examples). Our same cell line and chromosome cross-validation results are generated using hard pairs only. The cross-validation experiments were done in all autosomal chromosomes.

For the same cell line cross chromosome setting, we used the five random forests regression models trained on each fold from the training chromosome to predict contact counts for all pairs in a test chromosome. Each pair in the test chromosome had five predictions and we took the average of these predictions as the final predicted count. We note that in this setting, all test pairs are hard. Cross-chromosome experiments were done on five chromosomes, 9, 11, 14, 17 and 19.

For different cell line same chromosome setting, we again used the random forests regression models trained on the training data from each fold in one cell line and generated predictions for all pairs in the test cell line. Next, we took the average of these predictions as the final predicted count. When using MULTI-CELL features, we excluded the features derived from the regulatory signals measured in the test cell line. In this setting as well, all test pairs are hard pairs. Our ensemble approach took a further average of the predictions from all four training cell lines as the predictions in a test cell line. Cross-cell line experiments were done in three chromosomes: 14, 17 and 19.

### Feature extraction and representation

To extract features for HiC-Reg’s regression framework, we used data sets from the ENCODE project [42] for five cell lines: K562, Gm12878, Huvec, Nhek and Hmec, learning a separate model for each cell line. We selected 14 data sets that were measured in all five cell lines. These 14 data sets included ChIPseq datasets for 10 histone marks and CTCF, DNase I-seq and DNase I-seq derived motifs of RAD21 and TBP, which we had previously found to be helpful for predicting enhancer-promoter interactions in a classification setting [14]. A ChIP-seq signal is represented as the average read count aggregated into a 5kb non-overlapping bin. We obtained the raw fastq files from the ENCODE consortium [42], aligned reads to the human hg19 assembly using bowtie2 [43], retrieved reads aligned to a locus using SAMtools [44] and applied BEDTools [45] to obtain a base pair level read count. We next aggregated the read counts of each base pair in a 5kb region. Next, we normalized aggregated signal by sequencing depth and collapsed replicates by taking the median. As TBP and RAD21 ChIP-seq data are not available in Huvec, Nhek and Hmec cell lines, we predicted the binding sites using PIQ [46] on the Dnase I data and used the sum of purity scores for all motifs mapped to the same 5kb bin as the signal value.

We represented features of a region as a 14 dimensional feature vector, each dimension corresponding to one of the 14 genome-wide data sets. To generate a feature vector for a pair of regions, we used different strategies: PAIR-CONCAT, WINDOW and MULTI-CELL. In the PAIR-CONCAT case, we concatenated the 14-dimensional feature vectors of the two regions to obtain a feature vector of size 28. In the WINDOW case, we concatenated the 14-dimensional feature vectors of the two regions together with the feature vectors of the intervening region between the two regions to obtain a feature vector of size 42. We call this the WINDOW feature following Whalen et al [15]. The feature associated with the intervening region is a mean signal value of the feature in the region. In the MULTI-CELL case, we concatenated the 42-dimensional feature vectors of the two regions from all five cell lines to obtain a merged feature vector of size 210. Finally, for all these feature representations, we included genomic distance between the two regions of a pair as an additional feature.

### Application of Fit-Hi-C to call significant interactions

The output of HiC-Reg can be analyzed using a peak calling method such as Fit-Hi-C [22]. Fit-Hi-C uses spline models to estimate expected contact probability at a given distance. The input to Fit-Hi-C is a raw count interaction file and an optional bias file calculated by the ICE method [47]. Fit-Hi-C estimates the statistical significance of interactions using a binomial distribution and corrects for multiple testing using the Benjamini-Hochberg method and outputs the P-value and corrected Q-value for each pair of interactions. We adopted the two-phase spline fitting procedure and used a FDR~0.05 to define significant pairs. As our predictions are based on normalized contact counts, we directly used our predicted counts as input for Fit-Hi-C without bias file. For each cell line, we first concatenated predictions across all chromosomes and conducted Fit-Hi-C analysis on these pairs. We applied the same procedure on true counts to find significant pairs.

### Evaluation metrics

We used different metrics for assessing the quality of our predictions for contact counts. The distance-stratified Pearson’s correlation was used to directly measure the accuracy of the predicted counts. The other metrics were useful to assess the quality of results after further downstream analysis of HiC-Reg predictions and compare them to similar results obtained from actual measured data. In particular, enrichment of CTCF bi-directional motifs and ChIA-PET datasets enabled us to study the quality of significant interactions identified from HiC-Reg predictions, while the TAD similarity enabled us to study the ability of HiC-Reg predictions to capture structural units of organization.

#### Distance stratified Pearson’s correlation

To assess the quality of count prediction of HiC-Reg, we used Pearson’s correlation of predicted contact counts and true contact counts as a function of genomic distance. We grouped pairs of regions based on their genomic distance and calculated the Pearson’s correlation of predicted and true contact counts for pairs that fall into each distance bin. We considered all pairs upto a distance of 1MB in distance bins of 5kb. To easily compare the performance between different methods, chromosomes and cell lines, we summarized the distance-stratified Pearson’s correlation curve into the Area Under the Curve (AUC). AUC was computed using the Trapezoidal rule and divided by number of distance bins, so that it ranged from 0 to 1. The higher the AUC, the better the performance.

#### Enrichment of bidirectional CTCF motifs on significant interactions

Cell line-specific DNase I-seq fastq files were retrieved from ENCODE (http://hgdownload.soe.ucsc.edu/goldenPath/hg19/encodeDCC/wgEncodeOpenChromDnase/) and further processed with PIQ [46] to obtain the coordinates and orientation of CTCF motifs. PIQ gives a score from 0.5-1, which is proportional to the true occurrence of the motif. We selected a threshold of 0.9 to identify high confidence CTCF motifs. We use R1 to denote the region with the smaller starting coordinate, and R2 to denote the region with the larger starting coordinate. Following Rao et al. [21], an interaction is labeled as having a convergent CTCF orientation if R1 contains CTCF motifs on the forward strand (“+” orientation) and R2 contains CTCF motifs on the reverse strand (“−” orientation). We only focus on pairs with at least one CTCF motif mapped to R1 and at least one CTCF motif mapped to R2. A pair can have one of the four configurations: (i) “++” configuration where both R1 and R2 have the motifs in the “+” orientation, (ii) “+-” configuration where R1 has CTCF motifs in“+” orientation and R2 has CTCF motifs in the “−” strand, (iii) “-+” configuration, where R1 is in the “−” orientation and R2 is in the “+”, (iv) “–”, where both R1 and R2 have motifs in the “−” orientation. We counted the total number of pairs with each of these configurations and compared this with the number of significant pairs called by Fit-Hi-C using the Hypergeometric test and fold enrichment. Briefly, assume we are testing the enrichment for the “+-” configuration. Let the total number of possible pairs in the background be *S*. Let *k* be the total number of significant interactions with any of the configurations of CTCF motifs, let *m* to be the total number of interactions with the “+-” configuration of CTCF motifs and *q* be the number of significant interactions with the “+-” configuration of CTCF motifs. Using Hypergeometric test, we test the probability of observing *q* or more interactions of *k* interactions to have the “+-” configuration, given that there are *m* out of *S* total interactions that have the “+-” configuration. Fold enrichment is computed as 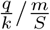.and must be greater than 1 to be considered as significant enrichment over background.

#### Validation of genome-wide predictions using ChIA-PET experiments

We downloaded 10 ChIA-PET data sets from Li et al. and Heidari et al.: PolII in HeLa and K562 and CTCF in the K562 cell line [23], and remaining seven ChIA-PET data sets from Heidari et al. [24], which included RNA PolII, CTCF, RAD21 and multiple chromatin marks in K562 and Gm12878 cell lines. Our metric for evaluating these genome-wide maps is fold enrichment, which assesses the fraction of significant interactions identified from HiC-Reg that overlapped with experimentally detected measurements, compared to the fraction of interactions expected by random chance. We mapped ChIA-PET interactions onto the pairs of regions used in HiC-Reg by requiring one region of an interaction from the ChIA-PET dataset to overlap with one region of a HiC-Reg pair (e.g., R1), and the other ChIA-PET region to map to the second region (e.g, R2). Fold enrichment is defined as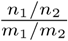, where *n*_1_ is the number of significant interactions from HiC-Reg that overlap with an interaction in the ChIA-PET data set, *n*_2_ is the total number of HiC-Reg significant interactions, *m*_1_ is the total number of interactions in the ChIA-PET dataset that can be mapped to any of the HiC-Reg pairs, and *m*_2_ is the total number of possible pairs in the universe. The observed overlap fraction of interactions is *n*_1_*/n*_2_ and the expected overlap fraction of interactions is *m*_1_*/m*_2_. A fold enrichment greater than 1 is needed in order to be considered significant.

#### Identification of TADs and TADs similarity

To identify topologically associated domains (TADs), we applied the Directionality Index (DI) method described in Dixon et al [32]. The method is based on Hidden Markov Model (HMM) segmentation of the DI. The DI is a score for a genomic region to measure the bias in the directionality of interactions for that region as measured in a Hi-C dataset. It is determined by the difference in the number of reads between the region and a genomic window (e.g., 2MB) upstream of the region and the number of reads between the region and a window downstream of the region. The window is user-defined. The DI score is segmented into three states of upstream, downstream or no bias. A TAD is then defined by a contiguous stretch of downstream biased states.

We transformed our upper triangle prediction matrix with resolution of 5kb to a symmetric interaction matrix and its genomic coordinates as the input. We used the default parameters of the package with a window size of 2Mb for defining the Directionality Index. For comparison, we applied the same procedure on SQRTVC normalized matrices (i.e. our true counts matrices) to identify TADs. We compared the similarity of TADs identified from the HiC-Reg predicted counts and true counts for each chromosome. Specifically, we matched a TAD found in the true count data to a TAD found in the predicted counts based on the highest Jaccard coefficient. The Jaccard coefficient measures the overlap between two sets, *A* and *B* and is defined as the ratio of the size of the intersection to the union.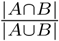 The Jaccard coefficient ranges from 0 to 1, with 1 denoting complete overlap. The Jaccard coefficients for each match were averaged across all TADs from the true counts. We repeated the matching procedure for each TAD from the predicted counts to a TAD in the true count and averaged the Jaccard coefficient across all TADs from the predicted counts. The overall similarity between TADs was then the average of these two averages.

### Feature Analysis

#### Single feature importance

To assess the importance of individual features we used Out of Bag Variable Importance measure [38], which computes importance of a feature based on the change in error on out of bag example pairs when the feature values are permuted. We used MATLAB’s implementation of this measure. In addition we counted the number of times a particular feature was used to predict the count for an example pair when it is part of the test set. Briefly, for each test example, *i* and each tree, *t* in the ensemble, let 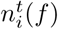denote the number of times feature *f* is used on the path from the root to the leaf in tree *t* for example *i*. The overall importance of feature *f* is 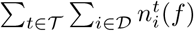 where, *T* stands for the ensemble of regression trees and *D* is the dataset of examples. We computed these counts on all test example pairs as well as examples with the top 5% lowest errors. The feature importances were very similar (**Supplementary Fig 7, 8**). A key difference between MATLAB’s OOB importance and our feature usage count importance is that the OOB is computed on left out examples from the training set, which includes the “easy” examples. The feature usage count is computed only on the hard pairs, which do not share a region with the training pairs.

#### Pairwise feature importance

To assess the importance of pairs of features, we considered all pairs of features that occur on the tree path from root to a leaf for a test example (similar to above). This approach is similar to the Foresight method [48], which counts the number of times a pair of features co-occur on the path from the root to the leaf. The main difference between our approach and that of Foresight is that we estimate these counts on the test examples, while Foresight estimates these on the examples in each leaf node in the tree identified during training. By using the counts on the test examples, our approach is less prone to overfitting. Because the feature rankings of individual features are similar when using all pairs and pairs with the top (smallest) 5% errors, we computed these counts only the top 5% error pairs.

#### Non-negative matrix factorization for identifying sets of features associated with sets of pairs

We developed a novel feature analysis method to identify feature sets associated with sets of pairs based on non-negative matrix factorization (NMF). The input to this approach is a*n* × *m* matrix *X* with *n* rows corresponding to test pairs and *m* columns corresponding to a pair of features and each entry *X*_*ij*_ corresponds to the number of times feature pair *j* is used to make a prediction for pair *i*. NMF decomposes the matrix *X* into two lower rank non-negative matrices *U* and *V*, where *X* = *UV* ^T^, *U* is *n* × *k, V* is *k* × *m*, and *k* is the rank. We used *k* = 5 for NMF. The *U* and *V* matrices are chosen to minimize squared error 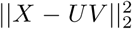 of the lower dimensional reconstruction. We use MATLAB’s non-negative matrix factorization function nnmf, with *k* = 5 factors to perform this factorization, which uses an iterative algorithm to estimate *U* and *V*. The *U* and *V* matrices provide a low-dimensional representation of the interaction pairs and feature pairs respectively. For ease of interpretation, we normalized the *U* matrix to make each row sum to unity (denoted as *Ū*) and normalized *V* matrix each column sum to unity (denoted as). To identify sets of features associated with sets of examples, we used the columns and rows of the *Ū*and 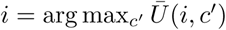 matrices. Specifically, we assigned a example *i* to cluster *c* if *i* = arg max_*c*_*I Ū* (*i, c′*). Similarly a feature pair, *j* is assigned to cluster *c* if 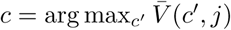. The columns of *U* and rows of *V* have one-to-one correspondence, thus providing a natural bi-clustering output. We were able to get striking factorization results where there were groups of pairs clearly associated with groups of pairs. To further interpret these feature pairs, we visualized the pairwise interactions as networks in Cytoscape [49], with node size proportional to the number of feature pairs they are associated with and edge weights corresponding to the strength of the association. The NMF based feature analysis enabled us to extract groups of examples associated with sets of feature pairs.

## Implementation and availability

The HiC-Reg code and associated MATLAB and R scripts to compute various validation metrics are available at https://github.com/Roy-lab/HiC-Reg/tree/master/Scripts.

## Acknowledgements

We thank the Center for High Throughput Computing at UW Madison for computational resources and Alireza Fotuhi Siahpirani for assistance with the CTCF motif and TAD analysis.

